# Sexually antagonistic coevolution between the sex chromosomes of *Drosophila melanogaster*

**DOI:** 10.1101/818146

**Authors:** Katrine K. Lund-Hansen, Colin Olito, Edward H. Morrow, Jessica K. Abbott

## Abstract

Antagonistic interactions between the sexes are important drivers of evolutionary divergence. Interlocus sexual conflict is generally described as a conflict between alleles at two interacting loci whose identity and genomic location are arbitrary. Here we build on previous theory and suggest that when these two loci are located on the sex chromosomes it can lead to cycles of antagonistic coevolution between them, and therefore between the sexes. We tested this hypothesis by performing experimental crosses using *Drosophila melanogaster* in which we reciprocally exchanged the sex chromosomes between five wild-type populations in a round-robin design. Disrupting putatively coevolved sex chromosome pairs resulted in increased male reproductive success in 16 out of 20 experimental populations (10 of which were significant), but also resulted in lower offspring egg-to-adult viability that affected both male and female fitness. After 25 generations of experimental evolution these sexually antagonistic fitness effects appeared to have been resolved. To help formalise our hypothesis, we developed population genetic models of antagonistic coevolution using fitness expressions based on our empirical results. Our models support the conclusion that antagonistic coevolution between the sex chromosomes is plausible under the fitness effects observed in our experiments. Together, our results lend both empirical and theoretical support to the idea that a cycle of antagonistic coevolution can occur between sex chromosomes and illustrates how this process may drive genetic and phenotypic divergence between populations.

**Significance:** Sex chromosomes are not only involved in genetic sex determination – they are also important factors in sexual conflict and speciation. Here, using a combination of experiments and population genetic models, we show that the sex chromosomes in *Drosophila melanogaster* can coevolve antagonistically. We found that swapping sex chromosomes between five *Drosophila melanogaster* populations increased male fitness at the cost of reduced offspring survival. After 25 generations, the increase had disappeared, consistent with the completion of a cycle of antagonistic coevolution. Using parameter values based on these empirical data, our models show that antagonistic coevolution between the sex chromosomes is a biologically plausible explanation for the results. Thus, our results point to a potentially important path to speciation through sexual conflict.

## Introduction

Sex chromosomes have a number of unique properties that distinguish them from autosomes, and one another, including their mode of inheritance, the selection experienced in the two sexes, and gene content (1). These processes are interdependent and strongly influence the genetic variation harboured on each sex chromosome. The Y chromosome is inherited exclusively through males, and therefore only exposed to selection in males. Because the Y chromosome is male-limited, any Y-linked genetic variation that is not male beneficial should experience purifying selection, which is consistent with empirical results (2–4). The X chromosome is inherited through males and females similarly to the autosomes, but experiences very different selection pressures than the autosomes or the Y (1). The X chromosome spends two-thirds of its time in females and is therefore largely exposed to selection in females (1), but since it is hemizygous in males, it is always exposed to selection when present in males (i.e., X-linked genes are unsheltered by dominance effects in males) (5). Thus, genes on the X chromosome are not sex-limited as on the Y chromosome, and whether they are female- or male-beneficial can depend on the dominance coefficient (6). Collectively, these unique properties have made sex chromosomes important factors in two major fields within evolutionary biology: speciation (7) and sexual conflict (1, 8).

Here we aimed to investigate whether interactions between sex chromosomes contribute to between-population divergence at the intra-specific level, as has previously been shown in interspecific comparisons (7). To do so, we performed experimental crosses to ‘swap’ either an X or a Y chromosome between five geographically isolated outbred populations of *D. melanogaster*, allowing us to isolate the effects of novel interactions between the sex chromosomes on male reproductive fitness. As we outline below, we expected to see different patterns of male reproductive success depending on whether population divergence is caused by incipient reproductive isolation, or is instead a result of sexually antagonistic coevolution between the sex chromosomes.

The accumulation of incipient ‘hybrid incompatibilities’ begins in allopatric populations within a species prior to speciation (9), and the X chromosome has been found to have a large effect on hybrid male sterility (10). So, if interactions between the sex chromosomes contribute to divergence between these populations primarily through accumulation of X-linked incompatibilities, we would expect to see a decrease in male fitness in the experimental populations. However, while previous studies have found evidence of early prezygotic isolation between sub-Saharan and cosmopolitan populations of *D. melanogaster* (11, 12), there is limited evidence for postzygotic isolation (13). So, we did not expect to find evidence of incipient reproductive isolation in our population crosses.

Sexually antagonistic coevolution between the sex chromosomes could, in principle, provide an alternative mechanism for the sex chromosomes to contribute to evolutionary divergence between populations (14), as suggested by studies of Y-linked regulation of gene expression (15). The theory of sexually antagonistic coevolution is based on the Red Queen-process (16). In short, when males increase their reproductive success through an adaptation that is simultaneously detrimental to females, it creates selection for a counter-adaptation in females to regain their lost fitness. The process may be repeated in multiple cycles over evolutionary time until a resolution is reached or a palliative adaptation ends the conflict (17). Sexually antagonistic coevolution can be considered a form of interlocus sexual conflict (14) if the traits involved are encoded by different genes in males and females (18). As first proposed by Rice & Holland in 1997, when the loci in question are located on the sex chromosomes, this could lead to cycles of sexual antagonistic coevolution between the sex chromosomes (14). If a Y-linked male-beneficial mutation arises that increases male reproductive success but decreases female fitness, it should spread through a population because it is subject to selection in males only. The sexually antagonistic male-beneficial mutation creates selection favouring compensatory mutations that might arise on the X chromosome or autosomes to restore female fitness again. Such mutations will likely differ between allopatric populations, and we would therefore expect to find an increase in male reproductive fitness when a Y chromosome with male beneficial mutations is paired with an X chromosome without the corresponding compensatory mutation(s). According to the sexually antagonistic coevolutionary model, we also predict a decrease in female fitness when mating with males harbouring a Y chromosome paired to a novel X chromosome.

Another corollary of the antagonistic coevolutionary model is that the effect of disrupting coevolved sex chromosomes on male and female fitness should decay over subsequent generations as new compensatory mutations arise, or novel combinations of segregating alleles achieve a similar compensatory effect. We tested this prediction by examining male and female fitness in our experimental populations 25 generations of experimental evolution. Crucially, the signature of antagonistic coevolution between the sex chromosomes in terms of male and female fitness differs from that for the accumulation of sex-linked incompatibilities, allowing us to identify whether one or other process is driving evolutionary divergence between populations.

To complement our empirical results and help formalize the hypothesis of sexually antagonistic coevolution between the sex chromosomes, we developed population genetic models describing the evolution of two interacting loci located in different genomic regions (i.e., unlinked): a Y-linked locus influencing both adult male fertilization success (i.e., sperm competition) and subsequent offspring survival, and a compensatory locus affecting only offspring survival located on either an autosome or the X chromosome.

## Results

### Empirical evidence for sexual antagonistic coevolution

To empirically test for evidence of population-specific interactions between the sex chromosomes in *D*. *melanogaster*, we crossed five outbred laboratory-adapted wild-type populations derived from four continents and three climatic zones in a round robin crossing design (Fig. 1). We created 20 novel populations where either the X chromosome (*novel X* treatment) or the Y chromosome (*novel Y* treatment) from one wild-type population (*wt*) was incorporated into another population (*SI Appendix A*, Fig. S1). We found a significant effect of treatment on male relative reproductive fitness (*SI Appendix A*, Table S1), with males from the *novel X* or *novel Y* treatments having a significantly higher relative fitness than *wt* males (Fig. 2*A*, *SI Appendix A*, Table S2). To exclude interactions between the novel sex chromosome and the autosomes as the cause, we also created a *novel XY* treatment where a pair of sex chromosomes from one *wt* population were introduced into another population (*SI Appendix A*, Fig. S2). We found no significant effect of treatment on relative fitness in the assay with *novel XY* treatments (Fig. 2*B*, *SI Appendix A*, Table S1). It therefore seems that introducing an individual novel sex chromosome caused the fitness differences rather than interactions with the autosomal background.

**Fig. 1:**
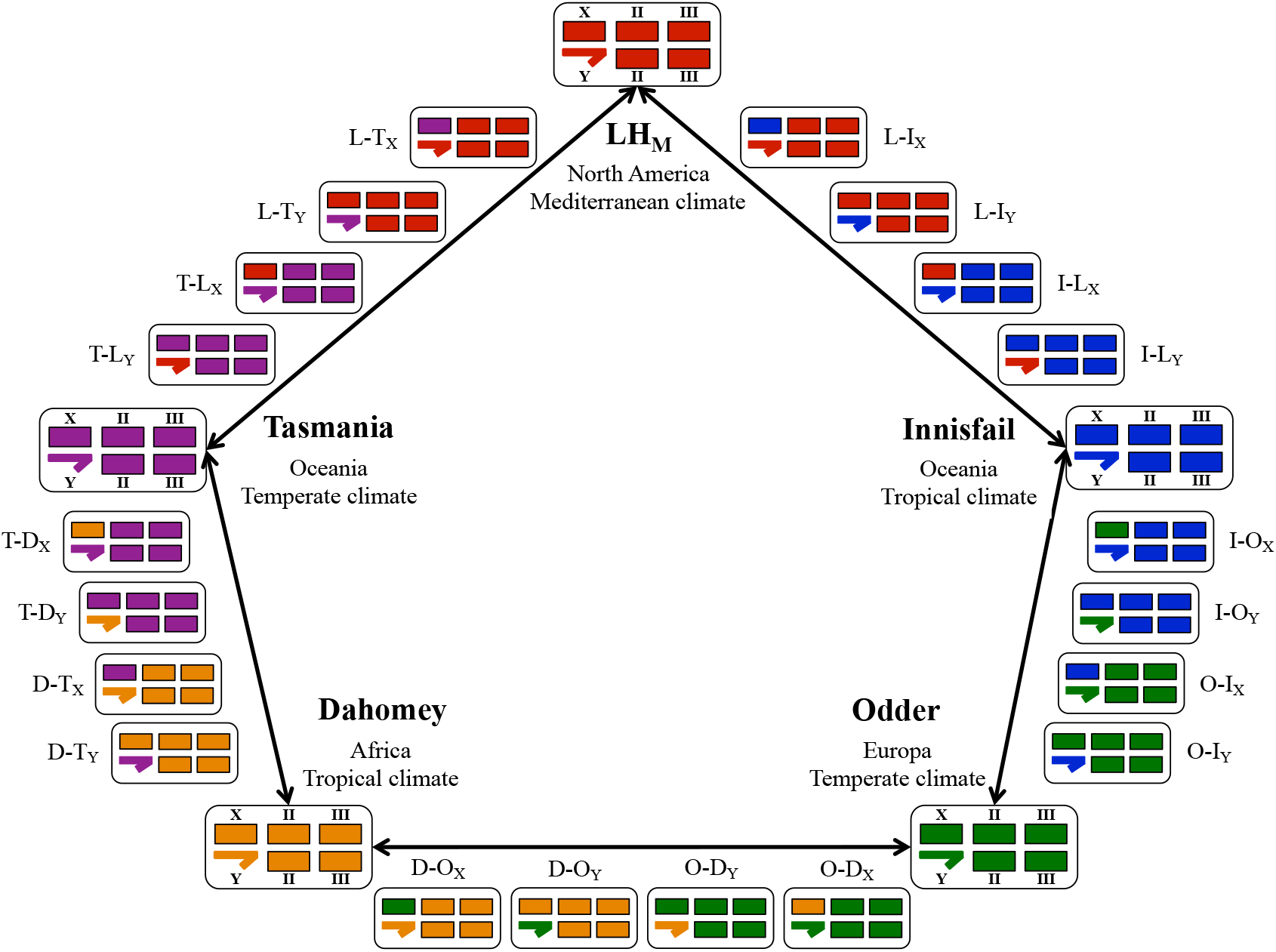
Schematic of the round robin experimental cross design. Each of the five wild-type populations is color-coded: LH_M_ (L, red), *Innisfail* (I, blue), *Odder* (O, green), *Dahomey* (D, orange), and *Tasmania* (T, purple). We crossed each wild-type population (large boxes) with two others from either a different continent or climate. This procedure generated four novel genotypes from each cross, so that every combination of the novel sex chromosome pairs experienced a different autosomal genetic background. In total, 10 *Novel X* and 10 *Novel Y* genotypes were generated (small boxes). Coloured bars represent the two major chromosomes (II and III) and chromosome X, with chromosome Y depicted by a half arrow. *Novel* genotypes are annotated as ‘genetic background – origin of sex chromosome’, with the novel sex chromosome indicated by an X or Y subscript.

**Fig. 2:**
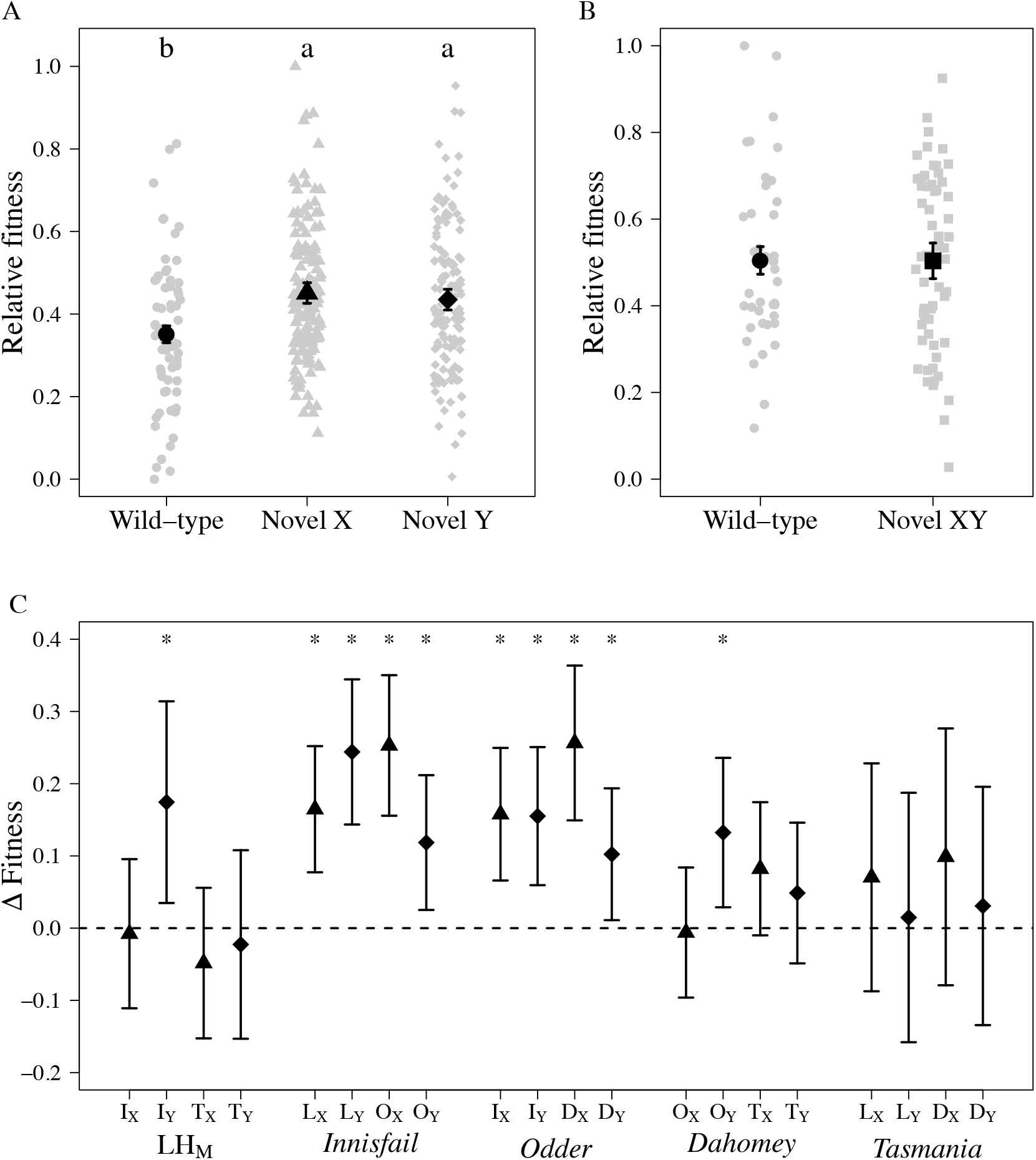
Male reproductive fitness assays at generation 0. (A) Relative reproductive fitness for wt (circles), novel X (triangle), and novel Y (diamonds) males. We detected statistically significant (P = 2.45e-04) increases in reproductive fitness for both novel X and novel Y males relative compared to the wild-type populations (Tukey HSD; P < 0.05). Mean (±SE) of fitted values from the linear model. The raw data is plotted as grey points. (B) Relative reproductive fitness for wt (circles) and novel XY (squares) treatment groups. There was no significant difference between wt and novel XY. Mean (±SE) of fitted values from the linear model. The raw data is plotted as grey points. (C) Change in relative fitness between the wt and the novel populations. ΔFitness = ωnovel population - ωwild-type) with bars indicating bootstrap 95% confidence (see Materials and Methods). 10 out of 20 novel populations (Novel X: triangle, novel Y: diamond) had significantly higher fitness than their wt counterpart (indicate by asterisks), and 16 populations had changed in a positive direction, which was more than expected by chance (P = 0.01). Each of the five wild-types are indicated underneath their four novel populations. Capital letters indicate which population the novel sex chromosome originates from (D: Dahomey, I: Innisfail, L: LH_M_, O: Odder, and T: Tasmania) and the novel sex chromosome is denoted by X or Y subscripts.

To investigate the interaction between the sex chromosomes in more detail, we calculated the difference in fitness between the novel populations and their *wt* counterparts (ΔFitness = **ω**_novel population_ - **ω**_wild-type_). We used bootstrap methods to model the data, and found that 10 of the 20 novel populations were significantly different from zero in a positive direction (Fig. 2*C*, *SI Appendix A,* Table S3). 16 out of the 20 point estimates were also positive, which is significantly different from what would be expected under random deviations (binomial test, *P* = 0.01).

### Male beneficial traits

To tease apart which fitness components were driving the overall pattern of increased relative fitness in the *novel* populations, we looked at a number of phenotypic traits that are correlated with male fitness. Interestingly, we found that the increase in fitness was related to different traits in the two *novel* sex chromosome treatments. For *novel X* males, we found that the increase in fitness was correlated with an increase in size, as *novel X* males were significantly larger than both *wt* and *novel Y* males (*SI Appendix A,* Fig. S3*A*, Table S1 & S2). For *novel Y* males, we found that the increase in fitness was correlated with an increased ability to displace other males’ sperm (*SI Appendix A*, Fig. S3*B*, Table S1), as *novel Y* males were significantly better at displacing sperm compared to *wt* males (*SI Appendix A,* Table S2).

### Female harmful traits

If the sex chromosomes coevolved antagonistically, the increase in male fitness should come at a cost to female fitness. We found a significant effect of treatment on total offspring number (*SI Appendix A*, Fig. S4*A*, Table S1), with *novel X* males siring a lower number of live offspring compared to *wt* males (Table S2). However, we did not find any difference in number of eggs laid by females mated with males from the different treatments (*SI Appendix A*, Fig. S4*B*, Table S1). The discrepancy between number of eggs and live offspring suggests a trade-off between offspring number and offspring quality, which was confirmed by a significant difference in egg-to-adult viability (SI Appendix A, Fig. S4*C*, Table S1). *Novel X* males sired significantly lower numbers of viable offspring than the other two treatments (*SI appendix A*, Table S2). Mating with *novel X* males decrease a female’s fitness by reducing the overall number of live offspring she produces. A decrease in offspring survival could be caused by meiotic drives seen in a departure from 50:50 sex ratio (19). We did not find a significant difference in sex ratio between the different treatments (*SI Appendix A*, Fig. S4*D*, Table S1).

We were unable to establish which harmful effects *novel Y* males had on female fitness through the assays performed in this experiment. But we were able to exclude costs to females associated with increased male harassment (e.g. due to larger size), males inducing high fecundity in females, and a reduction in offspring viability.

### Counter-adaptation

Any shift towards increased fitness in one sex at the cost of the other sex should lead to counter-adaption, which we expected to see as a reduction in male fitness over time. After 25 generations of experimental evolution (*SI Appendix A,* Fig. S5), we repeated both the male reproductive fitness assay and two of the significant phenotype assays to test if the interactions between the novel pairs of sex chromosomes had changed. As predicted, we no longer found a significant treatment effect on male fitness (Fig. 3*A*, *SI Appendix A,* Table S4). Indeed, we found that ΔFitness (ΔFitness = **ω**_novel population_ - **ω**_wild-type_) between novel populations and their wild-type counterparts had diminished in magnitude and were no longer significantly different from 0 (except for one population, I–O_X_) (Fig. 3*B*, *SI Appendix A,* Table S5). We also no longer found an effect of the treatments on sperm competition (*SI Appendix A,* Fig. S6*A*, Table S4) or offspring egg-adult survival (*SI Appendix A,* Fig. S6*B*, Table S4). There was a significant effect of the treatments on sex ratio (*SI Appendix A,* Fig. S6*C*, Table S4) as *novel Y* males produced significantly more male offspring than *wt* males (*SI Appendix A,* Table S6). However, this significant difference was not due to a change in sex ratio in the *novel Y* populations; rather it was due to a (putatively stochastic) change in the sex ratio in the *wt* populations.

**Fig. 3:**
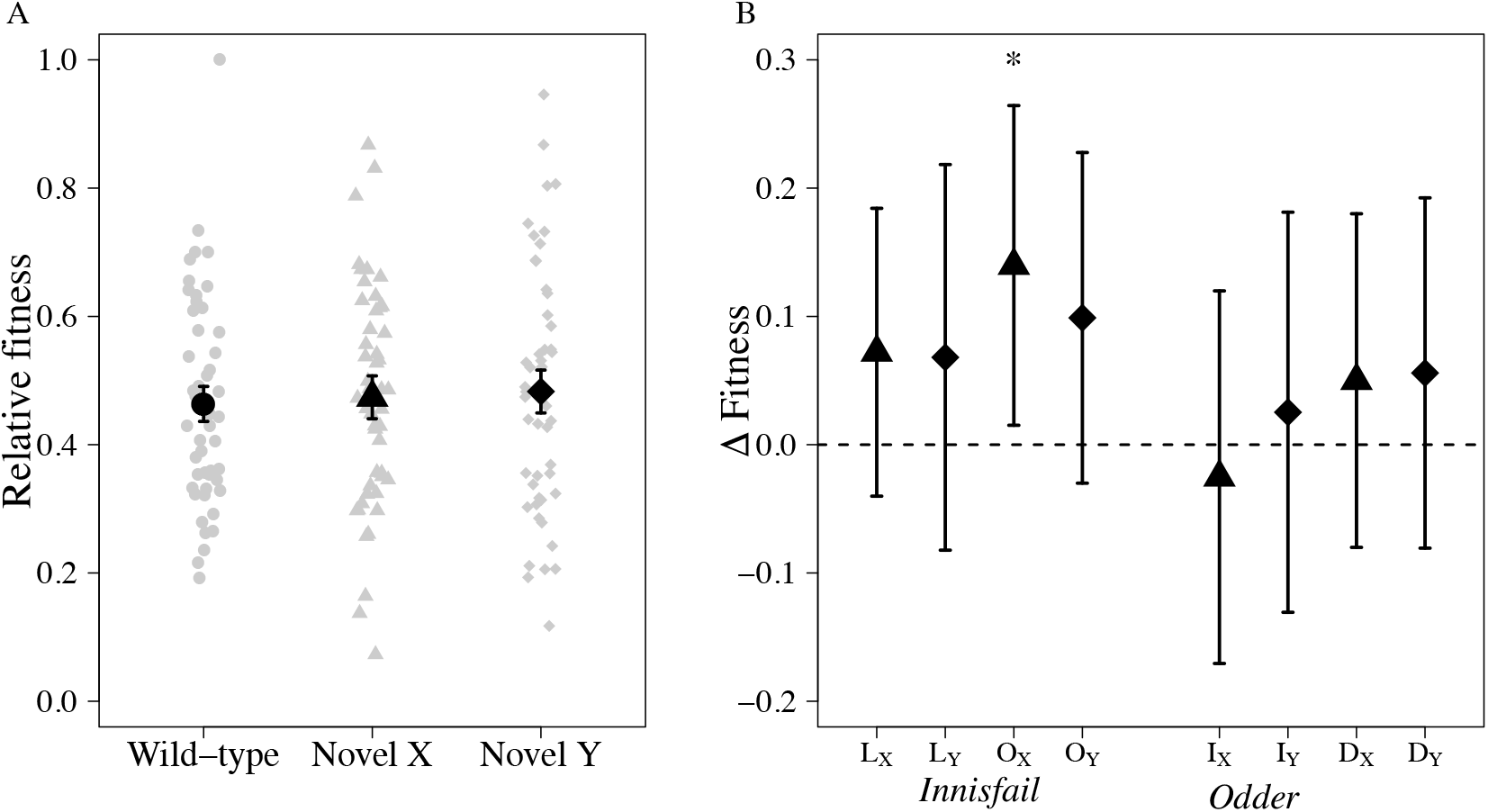
Reproductive fitness assays at generation 25. We selected 8 of the 10 significant populations for the evolution experiment. (A) Relative reproductive fitness for wt (black circle), novel X (triangle), and novel Y (diamonds) males. Novel treatment males no longer had a higher fitness relative to wt males after 25 generations of experimental evolution. Mean (±SE) of fitted values from the linear model. The raw data is plotted as grey points. (B) Change in relative fitness between wt and novel X (triangles) and novel Y (diamonds) populations (ΔFitness = ωnovel population - ωwild-type), with bootstrap 95% confidence intervals (see Materials and Methods). The asterisk indicates whether ΔFitness was significantly greater than zero (Tukey HSD; P < 0.05). Each of the wt populations are indicated underneath their four novel populations. Capital letters indicate which population the novel sex chromosome originates from (D: Dahomey, I: Innisfail, L: LHM, O: Odder) and the novel sex chromosome is denoted by X or Y subscripts.

### Population Genetic Models

Consider two alternative genetic systems involving two unlinked loci: a Y-linked locus (**Y**, with alleles *Y*, and *y*), and a compensatory locus located on either an autosome (**A**, with alleles *A*, and *a*) or the X chromosome (**X**, with alleles *X*, and *x*) (the Autosomal and X-linked models respectively), in a large population with discrete generations. In both models, a mutant *y* chromosome increases male fertilization success by a rate of 1 + *s*_*m*_ relative to the wild type (*Y*), but also reduces viability of offspring resulting from matings with females carrying the wild-type allele at the compensatory locus, *A* (or *X*). At the same time, females carrying the mutant *a* (or *x*) compensatory allele may incur a ‘cost of compensation’ in terms of offspring viability when mating with wild-type (*Y*) males (see Table 1 and Materials and Methods for a summary and description of fitness expressions). These fitness expressions create a scenario of antagonistic coevolution between a potentially male-beneficial Y-linked mutation, and the compensatory locus. Reminiscent of standard theories of compensatory evolution (e.g., 20, 21), each of the mutant alleles (*y* and either *a* or *x*) reduces offspring survival in isolation. However, unlike previous theory, the two loci are located on different chromosomes that do not recombine with one another; if a mutant *y* chromosome spreads to fixation, compensation at the population level requires that the compensatory mutation also fixes (see also recent models of “Father’s Curse” for similar scenarios, but without compensatory evolution or parental effects on offspring viability (22); a full description of the models is presented in the Materials and Methods and in *SI Appendix B*). Our theoretical analyses focus on (i) the evolutionary invasion of rare mutants at each locus individually; and (ii) single bouts of coevolution, beginning with invasion of a single-copy mutant *y* chromosome in a population initially fixed for the wild-type *Y* chromosome and for the wild-type *A* (or *X*) allele at the compensatory locus, and completing when both mutant alleles *y* and *a* (or *x*) become fixed (see 31 for a similar approach in the context of mito-nuclear coevolution).

**Table 1:**
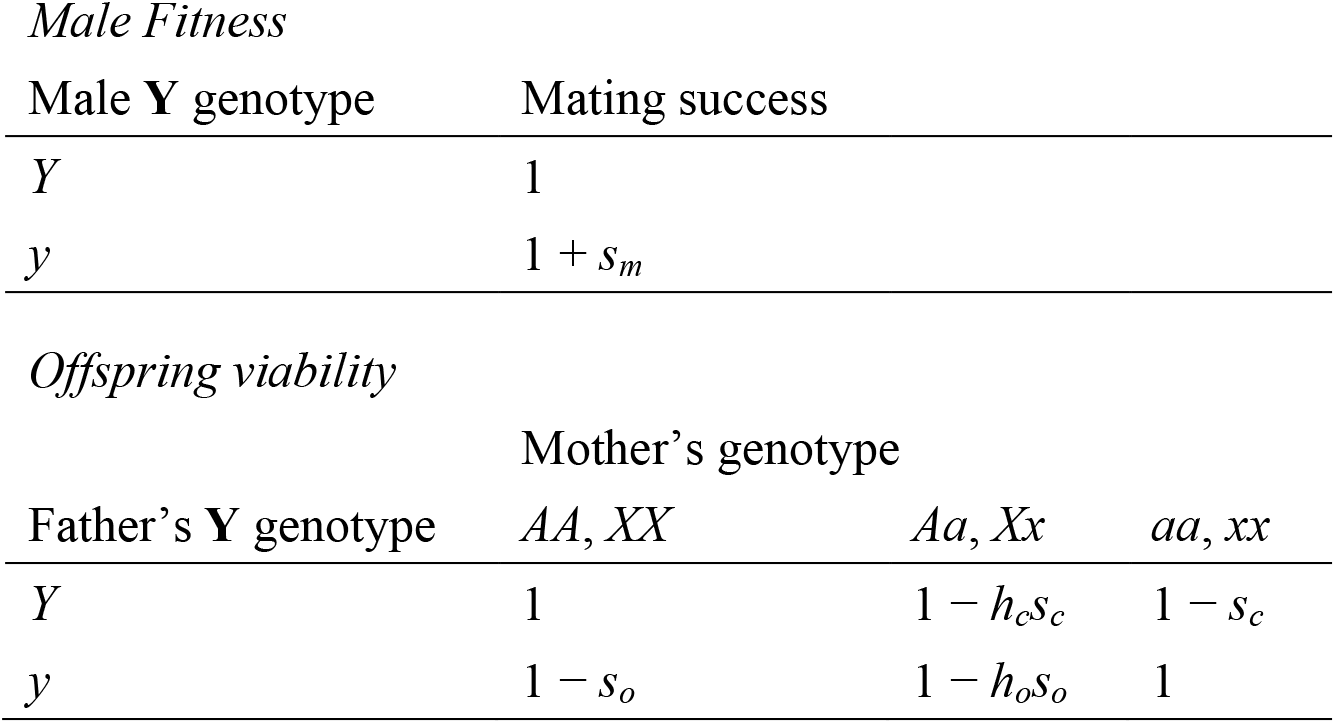
Fitness expressions for the Y-linked locus (**Y**) influencing male siring success, and the viability of offspring resulting from all possible combinations of parental genotypes at **Y** and the compensatory locus **A** (or **X**).

Evolutionary invasion analyses of the two models reveal tow key results. First, intuitively, invasion of a rare mutant *y* chromosome into a population initially fixed for the wild-type alleles at both loci requires that the increase in fertilization success for mutant males is greater than the accompanying reduction in offspring survival. Neglecting second order terms, a mutant *y* chromosome can spread when *δ* = *s*_*m*_ - *s*_*o*_ > 0. Second, whether the genomic location of the compensatory locus (either autosomal or X-linked) influences the invasion conditions for the compensatory mutation depends on the relative rate of mutation at the compensatory and Y-linked loci. If compensatory evolution is slow relative to the evolution of the Y-linked locus (e.g., the compensatory mutation rate is much smaller than the mutation rate to a mutant *y*), the mutant *y* chromosome is most likely to fix before a new compensatory mutation occurs. In this case, the genomic location of the compensatory locus does not influence the invasion conditions for a compensatory mutation. All males carry the mutant *y*, and so compensatory mutations will spread if there is any selection against the wild-type allele at the compensatory locus (i.e., 0 < *h*_*o*_,*s*_*o*_ < 1 for both models). Differences between the models emerge when new compensatory mutations can arise while the mutant *y* chromosome is still segregating in the population. If there is any cost of compensation for females (i.e., 0 < *s*_*c*_), compensatory mutations will experience purifying selection while *q*_*y*_ is small because most males carry the wild-type *Y* chromosome. As *q*_*y*_ increases, more matings involve mutant *y* males and females homozygous for the wild-type allele at the compensatory locus. When *q*_*y*_ reaches a threshold frequency, 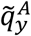 (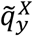 for the X-linked model), compensatory mutations will be favoured by selection (derivations provided in *SI Appendix B*). As illustrated in Fig. 4*A* for the case of additive compensatory fitness effects (*h* = *h*_*o*_ = *h*_*c*_ = 1/2), 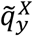 is always less than or equal to 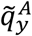, and the difference between the two thresholds is greatest when 0 < *s*_*o*_, *s*_*c*_ « *s*_*m*_. Overall, these results suggest that the genomic location of compensatory mutations (Autosomal or X-linked) will be most important when (i) the mutant *y* chromosome is strongly beneficial for males and there is little or no cost of compensation for females (i.e., when 0 < *s*_*o*_, *s*_*c*_ « *s*_*m*_); and (ii) when compensatory evolution is not limited by mutational variation (i.e., when mutant *y* chromosomes and compensatory mutations are likely to co-segregate in the population).

**Fig. 4:**
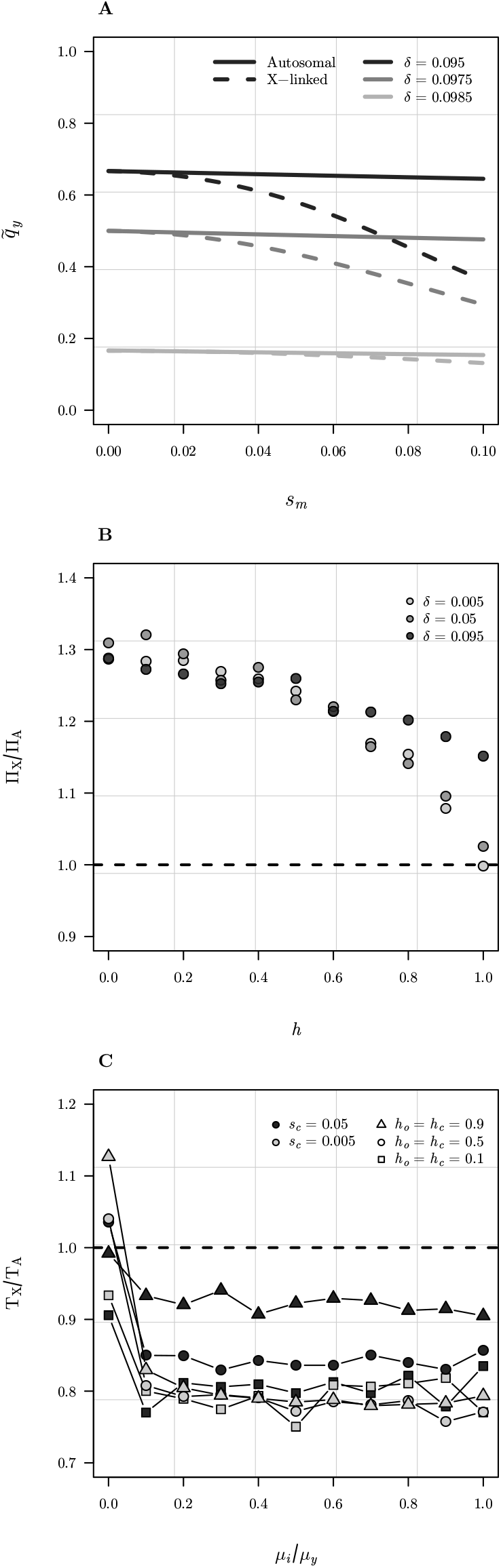
Summary of theoretical results. (A) Threshold frequencies of the mutant *y* chromosome at which a new compensatory mutation will experience positive selection for the Autosomal (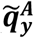, solid lines) and X-linked (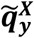, dashed lines) models respectively). As the mutant *y* chromosome sweeps to fixation, X-linked compensatory mutations will become selectively favoured earlier than autosomal ones. Results are shown for the case of additive fitness (*h*_*o*_ = *h*_*c*_ = 1/2), *s*_*m*_ = 0.1, and *s*_*c*_ = 0.005. (B) The relative probability of invasion for new Autosomal and X-linked compensatory mutations into populations initially fixed for the mutant *y* chromosome is always greater than or equal to 1, suggesting that when compensatory evolution is limited by mutational variation, the probability of invasion will be higher for X-linked compensatory mutations except under complete dominance. For simplicity we assumed equal dominance for compensatory fitness effects (i.e., *h* = *h*_*o*_ = *h*_*c*_). Results are shown for three different values of *δ* (recall that *δ* = *s*_*m*_ - *s*_*o*_), where *N* = 1,000, *s*_*m*_ = 0.1, and *s*_*c*_ = 0.005. Each point indicates the mean of 10^6^ replicate Wright-Fisher simulations for *δ* = 0.005 and *δ* = 0.05, but 5.0 × 10^6^ for *δ* = 0.095. (C) The relative time to complete a co-evolutionary cycle for the Autosomal vs. X-linked models plotted as a function of the relative mutation rates at the compensatory and Y-linked loci. To explore the consequences of slow vs. fast compensatory dynamics, *μ*_*Y*_ was held constant at 10^−3^, while *μ*_*i*_ (where *i* ∈ {***A,X***) was incremented from 10^−7^ to *μ*_*Y*_. At low values of *μ*_*i*_/*μ*_*Y*_, the rate of compensatory evolution is limited by mutational variation, but at intermediate to high values, the mutant *y* chromosome and compensatory mutations are more likely to segregate simultaneously. Results are shown for two different ‘costs of compensation’ (*s*_*c*_) and three different dominance scenarios (we again assume equal dominance, *h*_*o*_ = *h*_*c*_), and population size *N* = 1,000.

To complement our analytic results, we performed Wright-Fisher simulations to estimate two other important properties of coevolutionary cycles predicted by our models: (i) the probability of invasion of single-copy autosomal (Π_*A*_) and X-linked (Π_*X*_) compensatory mutations into populations of *N* individuals that are initially fixed for the mutant *y* chromosome; and (ii) the total time to complete a single bout of coevolution between the Y-linked and compensatory loci (*T*_*A*_ and *T*_*X*_, respectively) under recurrent mutation, selection, and drift. Although the invasion conditions for mutant compensatory mutations are the same for both models when the mutant *y* chromosome is initially fixed, the probability of eventual fixation for a single-copy mutation is larger for an X-linked compensatory locus (Π_X_) than an autosomal one (Π_A_) except for the special case of complete dominance (i.e., when *h*_*o*_ = 1) (Fig. 4*B*). The average time to complete a co-evolutionary cycle is also smaller for an X-linked than an autosomal compensatory locus, provided the compensatory mutation rate (*μ*_*a*_ or *μ*_*x*_) is not dramatically lower than the male-beneficial mutation rate (*μ*_*y*_) (Fig. 4*C*). When compensatory evolution is strongly limited by mutational variation, compensatory evolution can be faster on the autosomes than the X. This is due to an implicit trade-off between the probability of invasion, which is higher for the X than the autosomes (Fig. 4*B*), and the size of the mutational target, which is lower for the X because males only carry one copy.

## Discussion

We investigated whether if interactions between the sex chromosomes could contribute to between-population divergence at the intra-specific level. Taken together, our empirical results demonstrated that exchanging a sex chromosome between populations of *D. melanogaster* had an overall positive effect on male fitness. These results were consistent with the expected signature of antagonistic coevolution between the sex chromosomes on male fitness, but not with that of accumulated sex-linked incompatibilities influencing male sterility.

Mitochondria-nuclear interactions have previously been shown to affect male fertility in *D. melanogaster* (23), and both *novel X* and *novel Y* chromosomes were expressed with novel mitochondria. However, the *novel XY* chromosomes also were expressed with novel mitochondria and we found no effect on male fitness in these populations, suggesting that the magnitude of any mito-nuclear interactions was minor in these populations compared to the effect of interactions between the sex chromosomes.

We further wanted to know which specific phenotypic traits facilitated the increase in male reproductive fitness. We found that *novel X* males seemed to gain a mating advantage from an increase in size, as body size is an important factor in male fitness (24), and larger males have been shown to have a mating advantage (25). For *novel Y* males it seemed that they gained the advantage through being better at sperm displacement. Thus, the results suggest that different phenotypic and genetic mechanisms may be responsible for the increased male fitness observed in the two *novel* sex chromosome treatments.

Since the reproductive fitness results were consistent with antagonistic coevolution, we expected to find a corresponding decrease in female fitness. We found that offspring sired by *novel X* males had a significant lower egg-to-adult survival rate. We did not find the decrease in offspring survival to be caused by a distortion in sex ratio, which suggests the effect is not due to meiotic drive or sexually antagonistic zygotic drive (i.e. mortality resulting from competition between opposite-sex siblings (26). In addition, there are no known sex ratio meiotic drivers in *D. melanogaster*. Thus, the decrease in offspring survival was most likely caused by the father’s genotype. Reproduction is costly for females (27), so any eggs which do not produce live offspring are a expense and will over time reduce the reproductive fitness of the females. We were not able to identify which traits in females were negatively affected when mating with *novel Y* males. A possible cause to investigate in the future is the reduction of female receptivity after mating with males, which could lead to a reduction in female lifetime reproductive fitness.

As a final test of the hypothesis of antagonistic coevolution, we ran an evolution experiment for 25 generations to see if we would observe any counter-adaptation. After 25 generations, we found a reduction in fitness indicating strong selection pressure for compensatory evolution, which is supported by the simulation results. In principle, this subsequent reduction in male fitness could be due to new X-linked compensatory mutations; however, given the time-scale of the experiment it is more likely that any compensatory evolution by females must utilize standing genetic variation in the stock populations, with novel allelic combinations achieving a compensatory effect.

Collectively, our empirical results are consistent with predictions for antagonistic coevolution between the sex chromosomes. To further validate that we found evidence of antagonistic coevolution, we built a theoretical model based on our empirical evidence. Our theoretical results support the conclusion that antagonistic coevolution is (i) plausible under the fitness effects observed in our empirical experiments; and (ii) more likely to involve both sex chromosomes than the Y and an autosome, especially when compensatory dynamics are not slow relative to those of male-beneficial Y-linked mutations. An interesting corollary to these results is that antagonistic coevolution should drive more rapid among-population divergence on the X chromosome than the autosomes.

In conclusion, we empirically examined the role of the sex chromosomes in evolutionary divergence among allopatric populations of *D. melanogaster*. We found evidence of intragenomic conflict between the sex chromosomes that was independent of any interaction with the autosomes or mitochondria. Disrupting coevolved sex chromosomes resulted in an overall increase in reproductive fitness for *novel* males, through increased body size (for *novel X* males) and sperm displacement (for *novel Y* males), but also decreased offspring egg-to-adult viability. The accompanying reduction in offspring viability created indirect selection on females to mitigate the loss in fitness resulting from mating with a *novel* male. After 25 generations of experimental evolution *novel* males no longer enjoyed higher reproductive fitness and there were no differences in offspring survival, indicating that the sexually antagonistic effects of disrupting coevolved sex chromosomes had been resolved, probably by standing genetic variation for compensatory alleles in the experimental populations. Overall, our empirical results were consistent with a sexually antagonistic coevolutionary model of sex chromosome evolution in allopatric populations. Analysis of our theoretical models supports the plausibility of antagonistic coevolution between the sex chromosomes under the fitness effects observed in the experiments and predicts that such coadaptation is more likely to involve sex chromosomes than autosomes provided that compensatory evolution is not limited by mutational variation. These new insights into the interactions between the sex chromosomes can help further our understanding of early speciation events. Antagonistic coevolutionary cycles between the sex chromosomes will most likely follow different trajectories in different populations due to random mutations and the interaction between the environment and sexual conflict (28), resulting in genetic and phenotypic divergence between allopatric populations. Thus, over evolutionary time, antagonistic coevolutionary cycles could lead to hybrid incompatibility, and thereby contribute to speciation.

## Materials and Methods

### Empirical methods

#### *Drosophila* husbandry and fly stocks

We used five outbred laboratory-adapted wild-type populations: (1) *Dahomey*, Africa, tropical (25); (2) *Innisfail*, Oceania, tropical; (3) LH_M_, North America, mediterranean; (4) *Odder*, Europa, temperate (29); (5) *Tasmania*, Oceania, temperate. We maintained all wild-type populations under the standard LH_M_ culturing protocol (25°C, 12-12 light-dark cycle, 60% relative humidity, cornmeal-molasses-yeast medium, *SI Appendix A,* 30) for at least two generations before the cross. For the fitness assays we used an outbred LH_M_ population homozygous for the visible brown eye (*bw*) genetic marker (LH_M_-*bw*).

#### Male reproductive fitness

We estimated male reproductive fitness as the proportion of live offspring sired by males of the target treatment. Five adult target males were placed in a vial for two days with ten competitor LH_M_-*bw* males and 15 virgin LH_M_-*bw* females. The females were transferred into a single test tube to oviposit for 18 hours after which the females were discarded, and the test tubes left under standard LH_M_ conditions for 12 days. After which we counted the adult offspring and recorded their eye-colour to assess paternity. The *bw* genetic marker is recessive to the wild-type red eye-colour allele, so all red-eyed offspring can be assigned to red-eyed target males. We calculated relative fitness of the target males by dividing the fitness for each replicate by the maximum fitness across all replicates. We measured male fitness for three different assays: at generation 0 (*n* = 2 blocks × 7 experimental replicates × 25 populations; fitness estimates for 70 individuals per population), at generation 25 (*n* = 2 blocks × 3 experimental replicates × 2 replicate populations × 14 populations; fitness estimates for 60 individuals per population) and for the sex chromosome-autosome interactions (*n* = 10 experimental replicates × 10 populations; fitness estimates for 50 individuals per population).

Construction of the *novel* populations, the evolution experiment, male thorax size, sperm competition assay, male effect on female fecundity, and offspring egg-to-adult viability assay are described in *SI Appendix A*.

#### Statistical procedures

All the statistical analyses were conducted in R version 3.4.4 (31). We fit linear models with treatment as a fixed factor to test each of the dependent variables: thorax size and male effect on female fecundity, and tested for significant effects using ANOVA. We performed post-hoc Tukey HSD comparisons for all analyses with a significant treatment effect. We used linear mixed models (**lme4** 32) with treatment as a fixed factor and experimental block as a random factor to test the dependent variables: male reproductive fitness, egg-to-adult offspring viability, sex ratio, sperm competition, and total offspring number, and again tested for significant effects using ANOVA. We performed post-hoc Tukey HSD (**multcomp** 33) comparisons for all analyses with a significant treatment effect. We calculated bootstrap 95% confidence intervals around ΔFitness (ΔFitness = ω_novel population_ - ω_wild-type_) randomly resampling 13 out of 14 data points and recalculating ΔFitness 10,000 times. We used the Exact Binomial Test to test if there were more positive ΔFitness estimates than expected by chance. For statistical analysis after the 25 generations of experimental evolution we fit linear mixed models with treatment nested within replicate populations to test the dependent variables: male reproductive fitness, egg-to-adult offspring survival, sperm competition and sex ratio. For assays done in experimental blocks, we added experimental block as a random factor, tested for significant effects using ANOVA, and performed post-hoc Tukey HSD tests for analyses with significant treatment effect. We calculated bootstrap 95% confidence intervals around treatment means for ΔFitness by randomly resampling 11 out of 12 data points and recalculating ΔFitness 10,000 times.

### Theoretical models

We developed two population genetic models, identified by the location of the compensatory locus: the Autosomal and X-linked models respectively. We assumed discrete generations, and a life-cycle that proceeds as follows: (i) birth, (ii) selection on offspring survival, (iii) meiosis and mutation, (iv) selection on male mating success. Both loci in each model are biallelic. The Y-linked locus, **Y**, has wild-type *Y* and mutant *y* alleles with frequencies *q*_*Y*_ and *q*_*y*_ = 1 - *q*_*Y*_. The compensatory locus is denoted **A** (with alleles *A* and *a* and frequencies *q*_*A*_ and *q*_*a*_ = 1 - *q*_*A*_) for the autosomal model, and **X** (with alleles *X* and *x* and frequencies *q*_*X*_ and *q*_*x*_ = 1 - *q*_*X*_) for the X-linked model (wild-type alleles are indicated by capital letters, mutant alleles in lowercase). Following standard population genetics theory, **Y** is effectively haploid with strictly paternal inheritance, while the autosomal or X-linked compensatory loci (**A** or **X**) are diploid with bi-parental inheritance.

We based the fitness expressions used in our models loosely on our experimental results. The mutant *y* chromosome increases male mating success by a factor of 1 + *s*_*m*_ relative to the ancestral *Y* chromosome. Offspring survival depends on the paternal genotype at **Y** and the maternal genotype at the compensatory locus (**A** or **X**), as sires effects offspring viability (34) and any such negative effects should create selection for females to compensate by maternal effects. Offspring sired by mutant *y* males may experience reduced survival, depending on their mother’s genotype at the compensatory locus: matings involving parental genotypes [*y* : *AA*], [*y* : *Aa*], and [*y* : *aa*] result in offspring relative fitness of 1 - *s*_*o*_, 1 - *h*_*o*_*s*_*o*_, and 1 respectively. To make the models more general, and analytically tractable, we allowed females carrying the mutant *a* allele to incur a ‘cost of compensation’ when mating with wild-type (*Y*) males: matings involving parental genotypes [*Y* : *AA*], [*Y* : *Aa*], and [*Y* : *aa*] result in relative offspring fitnesses of 1, 1 - *h*_*c*_*s*_*c*_, and 1 - *s*_*c*_ respectively (Table 1). Similar to standard theories of compensatory evolution (e.g. 20, 21) each of the mutant alleles (*y* and either *a* or *x*) is deleterious in isolation. In our models, however, compensation requires that both parents have the appropriate mutant genotype at the other locus.

We model evolutionary invasion and single co-evolutionary cycles between the male-beneficial Y-linked locus and the compensatory locus (see 35 for a similar approach in the context of mito-nuclear coevolution). A bout of coevolution begins with the invasion of a single-copy mutant *y* chromosome in a population initially fixed for the wild-type *A* (or *X*) allele at the compensatory locus. The mutant *y* evolves under net positive selection if the increase in male mating success outweighs the reduction in offspring survival (i.e., if *δ* = (*s*_*m*_ - *s*_*o*_) > 0) until it becomes fixed (*q*_*y*_ = 1) or is lost from the population (*q*_*y*_ = 0). The compensatory locus (**A** or **X**) evolves under recurrent mutation and selection, with the population initially fixed for the wild-type *A* allele (*q*_*a*_ = 1 for the autosomal model) or *X* (*q*_*x*_ = 1 for the X-linked model). For simplicity we assume one-way mutation from *A* → *a* at a rate *u*_*a*_, and *X* → *x* at a rate *u*_*x*_ per meiosis. If females experience a ‘cost of compensation’ (i.e., when *s*_*c*_ > 0), the mutant compensatory locus will evolve under purifying selection until the mutant *y* chromosome reaches a threshold frequency, denoted 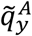 and 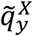, at which *y* becomes selectively favoured. A co-evolutionary cycle completes when the mutant allele becomes fixed at both loci (i.e., *q*_*y*_ = *q*_*a*_ = 1 or *q*_*y*_ = *q*_*x*_ = 1).

Our analytic results all assume large population sizes (negligible drift), and an equal sex ratio. To identify the conditions under which rare mutant alleles at each locus can spread during key points of an coevolutionary cycle, we performed a linear stability analysis for each model under three different scenarios: (i) invasion of mutant a *y* chromosome into populations initially fixed for the wild-type allele at both loci (initial frequencies of *q*_*y*_ = 0, and *q*_*i*_ = 0; where *i* ∈, {*a,x*}); (ii) invasion of a rare mutant compensatory allele into a population fixed for *y* (*q*_*y*_ = 1, and *q*_*i*_ = 0); and (iii) invasion of a mutant compensatory allele into a population with an arbitrary initial frequency of *y* (*q*_*y*_ = *q*_*y*_, *q*_*i*_ = 0). Mutant alleles can invade (i.e., the initial equilibrium is unstable) when the leading eigenvalue of the Jacobian of the system of recursions is greater than one (*λ*_*L*_ > 1; 43). When the initial frequency of *y* is arbitrary, solving the expression *λ*_*L*_ > 1 for *q*_*y*_ yields the threshold frequency at which compensatory mutations become selectively favoured, (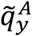 or 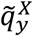). A full derivation of all models and analytic results is provided in *SI Appendix B*.

We complemented our analytic results with stochastic Wright-Fisher simulations for the Autosomal and X-linked models with population size *N* and an equal sex ratio. We estimated two important properties of co-evolutionary cycles from the simulations: (i) the probability of invasion of single copy autosomal (Π_A_) and X-linked (Π_X_) compensatory mutations into populations initially fixed for the mutant *y* chromosome; and (ii) the total time required to complete a single bout of coevolution between the Y-linked and compensatory loci (*T*_*A*_ and *T*_*X*_ for the autosomal and X-linked models respectively) under recurrent mutation, selection, and drift. Additional details and R code for the simulations can be found in *SI Appendix B*, and online at https://github.com/colin-olito/sexChromCoAdapt.

## Acknowledgments

Ary Hoffmann, Mads Fristrup Schou, and Stuart Wigby for the wild type fly stocks. Claire Webster and Qinyang Lee for technical support. Financial support came from Carl Tryggers Stiftelse (CTS 17:1) to K.K.L.-H., the European Research Council to J.K.A. (ERC-2015-StG-678148) and E.H.M. (ERC-2011-StG_20101109), a Royal Society University Research Fellowship to E.H.M, the Swedish Research Council (2011-3701) to E.H.M. and K.K.L.-H., and a Wenner-Gren International Postdoctoral Fellowship to C.O·.

## Notes

https://github.com/colin-olito/sexChromCoAdapt

