## Appendix A for "Sexually antagonistic coevolution between the sex chromosomes of *Drosophila melanogaster*" (from another version of this preprint: version 1, 2019-10-25)

### Supplementary material, Appendix A: Sexually antagonistic coevolution between the sex chromosomes of *Drosophila melanogaster*

Katrine K. Lund-Hansen[1,2*], Colin Olito[1], Edward H. Morrow[2], & Jessica K. Abbott[1]

[1] Department of Biology, Section for Evolutionary Ecology, Lund University, Lund 223 62, Sweden.
[2] School of Life Sciences, University of Sussex, Brighton BN1 9QG, United Kingdom

#### Materials and Methods

***Drosophila* husbandry and fly stocks**. All fly stocks were kept on a two-week regime with non-overlapping generations. On day 12 after oviposition, 16 pairs of adult flies were randomly selected from the pool of adults and placed in a food-vial with 6 mg yeast. The pairs were left in the vials for two days, and then flipped (transferred without $CO_2$) into fresh vials without yeast. The females were allowed to oviposit for 18 hours, after which all adult flies were discarded. The number of eggs was reduced to 150-200 per vial to maintain constant larval density.

For the creation of the novel populations we used clone-generator (CG) females with a double-X-chromosome (DX), a Y-chromosome, and two translocations of chromosome II and III (C(1)DX, *y*, *f*; T(2;3) *rdgC st in ri* $p^P$ $bw^D$).

**Construction of *novel X* and *Y* populations.** The five wild-type stocks were crossed in a round robin-scheme where one stock was crossed to two others. Pairings were performed reciprocally, generating four novel populations for each wild-type population, giving a total of ten *novel X* (X chromosome from a different population than the rest of the genome) and ten *novel Y* (Y chromosome from a different population than the rest of the genome) populations. The crosses were synchronised such that male fitness could be assayed simultaneously from all 20 novel populations.

To illustrate the principle of the entire crossing scheme we will employ the use of two hypothetical wild-type populations A and B:

*Novel Y* - Cross 1: CG females were crossed to A males (Fig. S1.1). This cross was performed with 672 breeding adults (♀32:16♂ per vial) in 14 vials and flipped once without reducing egg numbers. Cross 2: Heterozygote females (DX-T(2;3)/A) were crossed to *wt* A males a second time to produce the DX-A females required for crosses 3 and 4 in the *novel X* crossing scheme (see below; Fig. S1.2). This cross was performed with 640 breeding adults (♀16:16♂ per vial) in 20 vials and flipped twice without reducing egg numbers. Cross 3: Heterozygote females (DX-T(2;3)/A) were crossed to B males (Fig. S1.3), with 320 breeding adults (♀16:16♂ per vial) in ten vials and flipped twice without reducing egg numbers. Cross 4: B females were crossed to heterozygote males (T(2;3)/B) bearing a Y chromosome from A (Fig. S1.4) with 224 breeding adults in seven vials without reducing egg numbers. This creates *novel Y* males, for use in subsequent assays, where Y chromosomes from A have been placed in a B genetic background (Fig. S1.5).

*Novel X* - Cross I: CG females were crossed to B males (Fig. S1.I), with 672 breeding adults (♀32:16♂ per vial) in 14 vials and flipped once without reducing egg numbers. Cross II: Heterozygote females (DX-T(2;3)/B) were crossed to B males (Fig. S1.II), with 640 breeding adults (♀16:16♂ per vial) in 20 vials and flipped twice without reducing egg numbers. Cross III: DX-A females (from cross 2 in *novel Y*) were crossed to heterozygote males (T(2;3)/B) (Fig. S1.III), with 320 breeding adults (♀16:16♂ per vial) in ten vials and flipped twice without reducing egg numbers. Cross IV: DX-A females (from cross 2 in novel Y) were crossed to heterozygote males (T(2;3)/A) bearing a X chromosome from B (Fig. S1.IV). This cross was performed with 224 breeding adults (♀16:16♂ per vial) in seven vials without reducing egg numbers. This creates *novel X* males, for use in subsequent assays, where X chromosomes from population B have been placed into a population A genetic background (Fig. S1.V).

In both crossing schemes it should be noted that since there is no balancer for the small dot chromosome IV, we could not control how it was inherited. The dot chromosome only makes up 1% of the total genome and should thus have a limited effect (43). Since the dot chromosome is almost non-recombining (44), at the end of the *novel Y* crosses the probability of the dot chromosome being from the corresponding autosome population is 0.4. For the *novel X* crosses the probability is 0.5, and any effects of chromosome IV origin would be averaged out across the multiple individuals assayed.

**Construction of novel XY populations.** To estimate the effect of changing the autosomal background when both sex chromosomes were transferred together, we crossed four wild-type populations (*Dahomey*, *Innisfail*, $LH_M$, and *Odder*) to create novel populations where X and Y chromosomes from one population are placed into a genetic background of the two largest autosomes from a second population. As with the *novel X*/*Y* experiment, crosses were synchronized so that male fitness could be assayed simultaneously for all novel populations. Again, we illustrate the crossing scheme using two hypothetical populations A and B.

*Novel XY* - Cross 1: DX-A females were crossed to A males with a Y chromosome from B (Fig. S2.1), with 224 breeding adults (♀16:16♂ per vial) in 7 vials. Cross 2:

DX-A females with a Y chromosome from B were crossed to A males with a X chromosome from B (Fig. S2.2), with 448 breeding adults (♀16:16♂ per vial) in 14 vials. This creates males with both sex chromosomes from B but autosomes from A (Fig. S2.3). The probability of the dot chromosomes being from the same population background as autosome II and III was in this case 0.7.

**Evolution experiment protocol.** The three treatments (*novel X*, *novel Y*, and *wt*) were kept in an adult breeding population of 160 pairs (♀ 16:16 ♂ per vial) in 10 vials, with a minimum of 1,500 offspring. There were four populations of each treatment, with each population kept in two replicates.

*Novel X* – Males carrying a novel X chromosome were crossed at each generation to females who carried a double-bound X-chromosome (DX) and a Y chromosome (Fig. S5, *Novel X*). Because only half of the offspring of DX females survive to adulthood, the number of eggs were reduced to ~300 for these populations. The other half is inviable because they receive the wrong complement of sex chromosomes, either two Y chromosomes or three X chromosomes.

*Novel Y* – At each generation males carrying a novel Y chromosome were cross to X/X females (Fig. S5, *Novel Y*).

*Wt* population – The four *wt* populations were kept at the same population size as the novel treatments.

**Selected populations.** For all the following phenotypic assays we chose to assay novel populations with a ΔFitness significantly different from zero (Table S**3**). For the evolution experiment we selected four *novel X* populations (*Innisfail*–$L_X$, *Innisfail*–$O_X$, *Odder*–$I_X$, and *Odder*–$D_X$), four *novel Y* populations (*Innisfail*–$L_Y$, *Innisfail*–$O_Y$, *Odder*–$I_Y$, and *Odder*–$D_Y$), and their four corresponding *wt* populations (*Dahomey*, *Innisfail*, $LH_M$, and *Odder*).

**Male thorax size.** Body size was estimated using measurements of thorax length. On day 12 after oviposition, 50 males from the selected populations were placed in 95 % ethanol. Air-dried males were measured using a Nikon SMZ800 dissecting microscope at 63x magnification fitted with an eyepiece graticule. To ensure repeatability of the measurements, one person measured all the flies and we presumed that any bias would be equalised over the populations.

**Sperm competition assay**. Sperm offence (P2) was determined by the share of paternity achieved by the last male mated to a female. Five $LH_M$-*bw* virgin females were combined for an hour with ten $LH_M$-*bw* males. The females were the left on their own for 48 hours where after they were combined for 24 hours with ten target males from the selected populations. The females were then isolated in test tubes for 18 hours and the tubes were left under standard $LH_M$ conditions for 12 days. The adult offspring of the five females were counted and scored for eye-colour to assess paternity. P2 was assayed at two different time points: generation 6 (two blocks, seven experimental replicates per selected population in each) and generation 30 (one block with seven experimental replicates per selected population).

**Male effect on female fecundity**. Male effect on female fecundity was measured as the number of eggs laid by females during an 18 hour period, which corresponds to the period normally available to females from the base population (41). Five $LH_M$ virgin females were collected on the day of eclosion and combined with five males from the selected populations on day 12 after oviposition. After two days the five females were isolated into single test-tubes for 18 hours to lay eggs and were then discarded, while the test-tubes were frozen so the eggs could be counted at a later date. The fecundity for each female was averaged across the number of eggs laid by the five females. This assay was performed in nine experimental replicates per population.

**Offspring egg-to-adult viability assay**. Offspring egg-to-adult viability was estimated as the proportion of 100 eggs that developed into live adults within 12 days. 20 pairs of flies from each of the selected populations were placed in bottles with a Ribena-agar medium plate overnight to oviposit (Ribena is a blackcurrant-flavoured juice drink). 100 eggs were counted and transferred to a fresh food-vial, and left under standard $LH_M$ conditions. After 12 days all offspring and pupae were counted and sexed for estimation of sex ratio. Egg-to-adult viability was assayed at two different time points: generation 4 (two blocks, five experimental replicates per selected population in each) and generation 39 (four blocks with six experimental replicates per selected population per replicated population in one block).

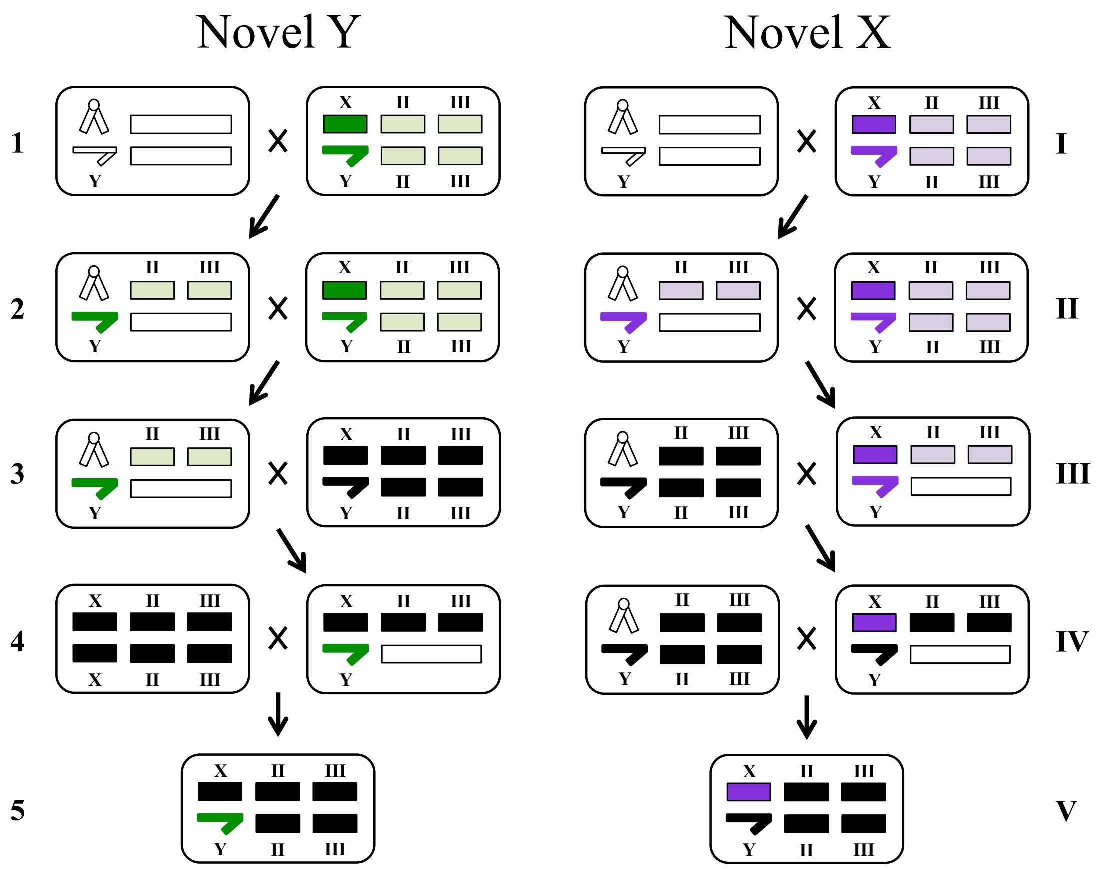


**Fig. S1:** Protocol for the crosses to create novel populations. Autosomes (II and III) and chromosome X are depicted as rectangles, and the autosome translocation (2:3) is depicted as the elongated white rectangle. The double-X-chromosome is depicted as Ѧ and the chromosome Y as a half arrow. Only offspring used for the cross in the next generation is shown. For additional details see *Construction of novel X and Y populations*.

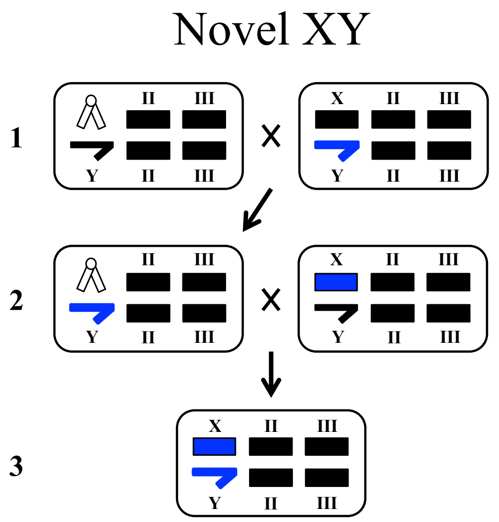


**Fig. S2:** Protocol for the crosses to create *novel XY* populations. Autosomes (II and III) and chromosome X are depicted as rectangles, the double-X-chromosome is depicted as Ѧ, and the chromosome Y as a half arrow. Only offspring used for the cross in the next generation is shown. For additional details see *Construction of novel XY populations*.

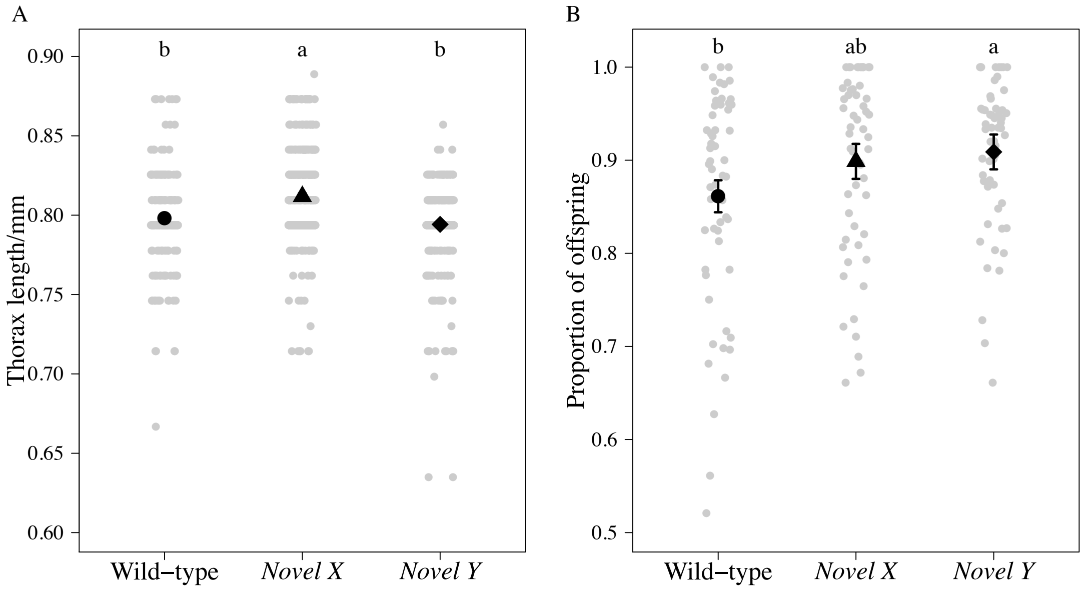


**Fig. S3:** Assays to investigate the benefits of the *novel* sex chromosomes Mean (±SE) of fitted values from the linear models. *wt:* black circle, *novel X:* triangle**,** *novel Y:* diamonds**.** The raw data is plotted as grey points. **(**A) Thorax size. *Novel X* males were significant larger than the two other treatments. Letters indicates significance (Tukey HSD; $P < 0.05$). (B) Sperm competition. *Novel Y* males were significantly better at replacing sperm than the other two treatments. Letters indicate significance (Tukey HSD; $P < 0.05$).

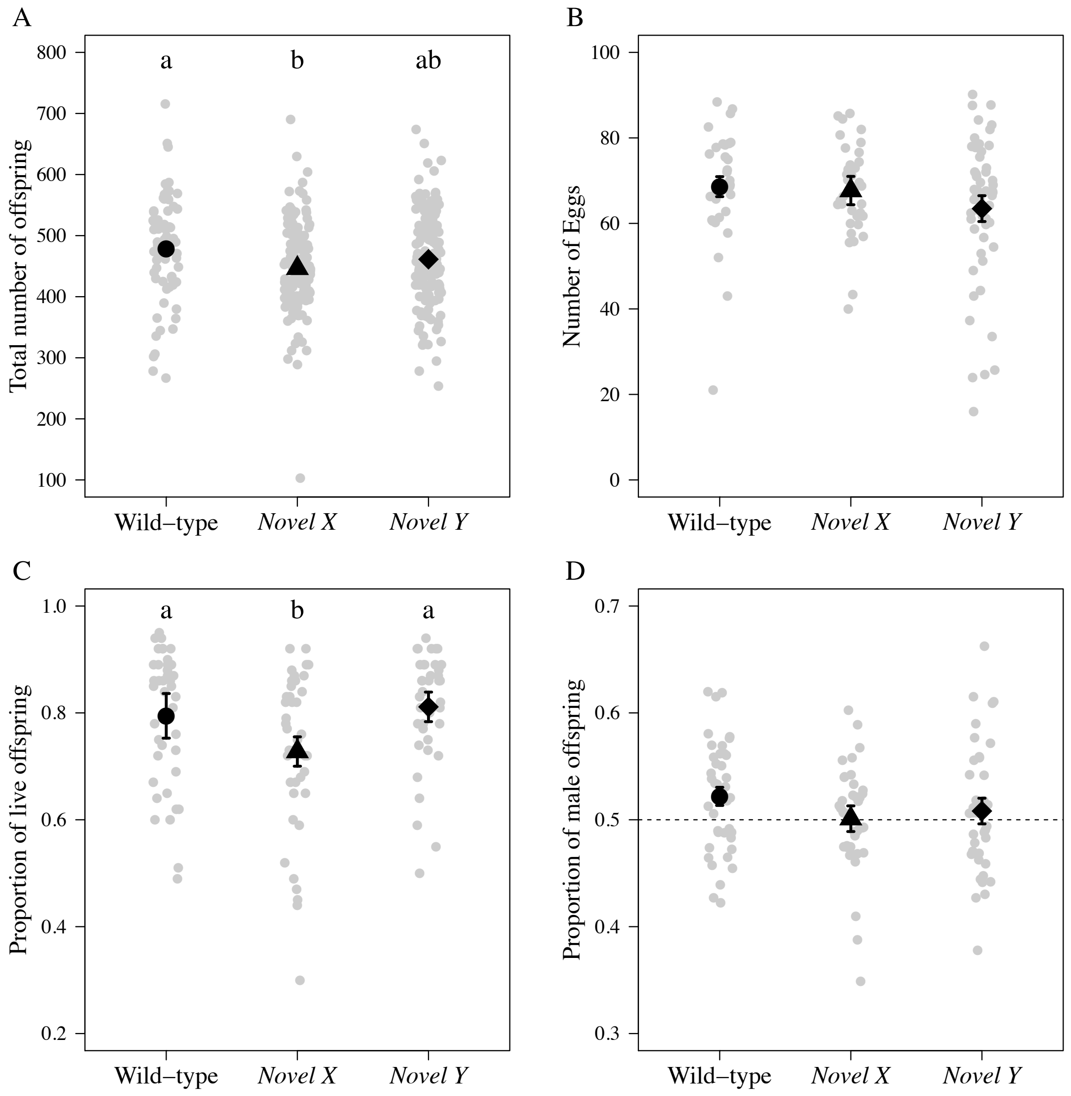


**Fig. S4:** Assays to investigate the cost of the *novel* sex chromosomes**.** Mean (±SE) of fitted values from the linear models. *wt:* black circle, *novel X:* triangle**,** *novel Y:* diamonds**.** The raw data is plotted as grey points. **(**A) Total number of offspring. The total number of offspring produced by $LH_M$-*bw* females in the male reproductive fitness assay. There was a significant lower number of offspring sired by *novel X* males. Letters indicates significance (Tukey HSD; $P < 0.05$). ) (B) Male effect on female fecundity. There was no effect of males from the novel treatments on female fecundity. (C) Egg-to-adult offspring viability. The offspring of *novel X* males had a significant lower survival than the two other treatments. Letters indicates significance (Tukey HSD; $P < 0.05$) (D) Sex ratio. There was no significant difference in the proportion of male offspring produced by *wt* and novel treatment males.

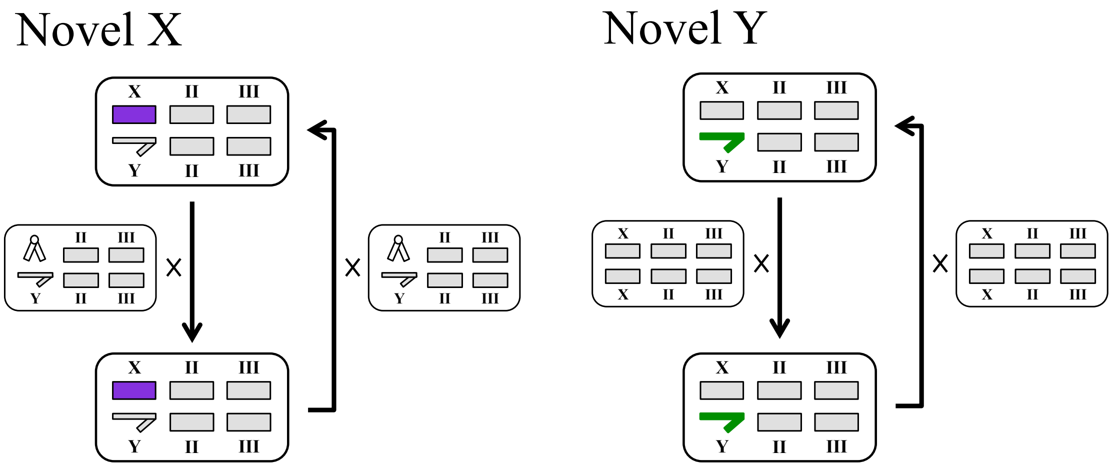


**Fig. S5:** The experimental setup for the evolution experiments for the novel populations. Autosomes (II and III) and chromosome X are depicted as rectangles, the double-X-chromosome is depicted as , and the chromosome Y as a half arrow. Only offspring used for the cross in the next generation is shown. For additional details see *Evolution experiment protocol*.

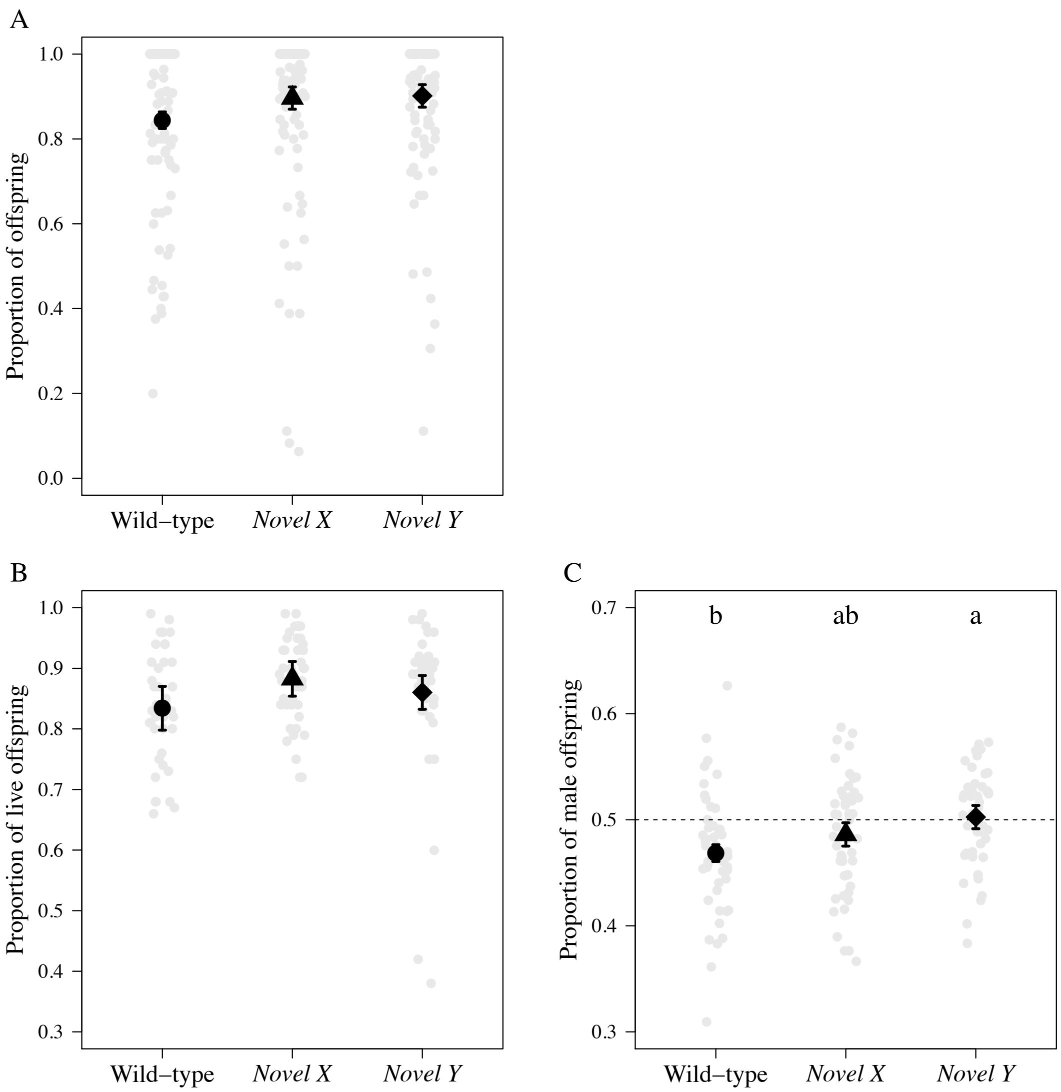


**Fig. S6:** Assays at the end of the evolution experiment. Mean (±SE) of fitted values from the linear models. *wt:* black circle, *novel X:* triangle**,** *novel Y:* diamonds**.** The raw data is plotted as grey points. **(**A) Sperm competition. After 32 generations there was no longer a significant effect of *novel Y* males ability to replace sperm compared to the other two treatments. (B) Egg-to-adult offspring viability. After 39 generations there was no longer a significant effect of *novel X* males on offspring survival. (C) Sex ratio. After 39 generations *novel Y* males produce significantly more male offspring than the *wt* treatment. Letters indicate significance (Tukey HSD; $P < 0.05$).

Table S1: Summary of the results from ANOVA analysis of fitting linear models (thorax size and male effect on female fecundity) and linear mixed-effects models (male reproductive fitness, egg-to-adult offspring viability, sex ratio, sperm competition, and total offspring number).

| Source | df | F | P |
|---|---|---|---|
| Relative male fitness, *Novel X & Y* | | | |
| Treatment groups | 2 | 8.52 | $2.44e^{-04}$ |
| Experimental block | 1 | | 1 |
| Relative male fitness, *novel XY* | | | |
| Treatment groups | 1 | $5e^{-04}$ | 0.98 |
| Thorax size | | | |
| Treatment groups | 2 | 61.01 | $<2.2e^{-16}$ |
| Sperm competition, P2 | | | |
| Treatment groups | 2 | 3.58 | *0.03* |
| Experimental block | 1 | | 0.34 |
| Total number of offspring | | | |
| Treatment groups | 2 | 4.10 | *0.02* |
| Experimental block | 1 | | 1 |
| Male induced female fecundity | | | |
| Treatment groups | 2 | 1.75 | 0.18 |
| Egg-to-adult viability | | | |
| Treatment groups | 2 | 5.05 | $7.92e^{-03}$ |
| Experimental block | 1 | | $5.17e^{-03}$ |
| Sex ratio | | | |
| Treatment groups | 2 | 1.56 | 0.21 |
| Experimental block | 1 | | 1 |

Table S2: Summary of the results from the post-hoc Tukey test of analyses with a significant treatment effect in the linear models and linear mixed-effects models.

| Source | P |
|---|---|
| Relative male fitness, *Novel X* & *Y* | |
| *Novel X – wt* | *< 0.001* |
| *Novel Y – wt* | *$2.31e^{-03}$* |
| *Novel X – Novel Y* | 0.71 |
| Thorax size | |
| *Novel X – wt* | *$3.46e^{-11}$* |
| *Novel Y – wt* | 0.13 |
| *Novel X – Novel Y* | *$<2.2e^{-16}$* |
| Sperm competition, P2 | |
| *Novel X – wt* | 0.12 |
| *Novel Y – wt* | *0.03* |
| *Novel X – Novel Y* | 0.85 |
| Total number of offspring | |
| *Novel X – wt* | *0.01* |
| *Novel Y – wt* | 0.28 |
| *Novel X – Novel Y* | 0.25 |
| Egg-to-adult viability | |
| *Novel X – wt* | *0.04* |
| *Novel Y – wt* | 0.83 |
| *Novel X – Novel Y* | *$8.00e^{-03}$* |

Table S3: The change in relative fitness with confidence intervals (CI) calculated by bootstrap for each of the novel populations. The populations marked in bold are significantly different from zero. Annotation for the novel populations is genetic background – origin of sex chromosome (X or Y). D: *Dahomey*, I: *Innisfail*, L: LH$_M$, O: *Odder*, and T: *Tasmania*.

| | ΔFitness | 95 % CI |
|---|---|---|
| Novel X | | |
| LH$_M$–I$_X$ | -0.0007 | -0.11; 0.10 |
| LH$_M$–T$_X$ | -0.05 | -0.15; 0.06 |
| ***Innisfail*–L$_X$** | **0.16** | **0.08; 0.25** |
| ***Innisfail*–O$_X$** | **0.27** | **0.15; 0.35** |
| ***Odder*–I$_X$** | **0.15** | **0.07; 0.25** |
| ***Odder*–D$_X$** | **0.27** | **0.15; 0.36** |
| *Dahomey*–O$_X$ | -0.008 | -0.10; 0.08 |
| *Dahomey*–T$_X$ | 0.04 | -0.01; 0.17 |
| *Tasmania*–L$_X$ | 0.07 | -0.09; 0.23 |
| *Tasmania*–D$_X$ | 0.09 | -0.08; 0.28 |
| Novel Y | | |
| **LH$_M$–I$_Y$** | **0.14** | **0.03; 0.31** |
| LH$_M$–T$_Y$ | -0.04 | -0.16; 0.11 |
| ***Innisfail*–L$_Y$** | **0.22** | **0.14; 0.34** |
| ***Innisfail*–O$_Y$** | **0.11** | **0.03; 0.21** |
| ***Odder*–I$_Y$** | **0.14** | **0.06; 0.25** |
| ***Odder*–D$_Y$** | **0.10** | **0.01; 0.19** |
| ***Dahomey*–O$_Y$** | **0.10** | **0.03; 0.23** |
| *Dahomey*–T$_Y$ | 0.02 | -0.05; 0.14 |
| *Tasmania*–L$_Y$ | 0.004 | -0.16; 0.19 |
| *Tasmania*–D$_Y$ | 0.02 | -0.14; 0.20 |

Table S4: Summary of the results from ANOVA analysis of linear mixed-effects models for the evolved populations.

| Source | df | F | P |
|---|---|---|---|
| Relative fitness for *Novel X & Y* | | | |
| Treatment groups | 2 | 0.17 | 0.85 |
| Nested factor | 1 | | 1 |
| Block | 1 | | $2.22e^{-06}$ |
| Sperm competition, P2 | | | |
| Treatment groups | 2 | 2.78 | 0.06 |
| Nested factor | 1 | | 1 |
| Egg-to-adult viability | | | |
| Treatment groups | 2 | 1.10 | 0.43 |
| Nested factor | 1 | | 0.25 |
| Block | 1 | | $7.37e^{-08}$ |
| Sex ratio | | | |
| Treatment groups | 2 | 4.77 | $9.92e^{-03}$ |
| Nested factor | 1 | | 0.25 |
| Block | 1 | | $7.37e^{-08}$ |

Table S5: The change in relative fitness between the novel evolved populations and the wild-type background with confidence intervals (CI) calculated by bootstrap. The population marked in bold is significantly different from zero. Annotation for the novel populations is genetic background – origin of sex chromosome (X or Y). D: *Dahomey*, I: *Innisfail*, L: $LH_M$, and O: *Odder*.

| | ΔFitness | 95 % CI |
|---|---|---|
| Novel X | | |
| *Innisfail*–$L_X$ | 0.07 | -0.04; 0.18 |
| ***Innisfail*–$O_X$** | **0.14** | **0.02; 0.26** |
| *Odder*–$I_X$ | -0.05 | -0.17; 0.12 |
| *Odder*–$D_X$ | 0.03 | -0.08; 0.18 |
| Novel Y | | |
| *Innisfail*–$L_Y$ | 0.09 | -0.08; 0.22 |
| *Innisfail*–$O_Y$ | 0.10 | -0.03; 0.23 |
| *Odder*–$I_Y$ | 0.02 | -0.13; 0.18 |
| *Odder*–$D_Y$ | 0.04 | -0.08; 0.19 |

Table S6: Summary of the results from the post-hoc Tukey test of analyses with a significant treatment effect in the linear mixed-effects models.

| Source | P |
|---|---|
| Sex ratio | |
| *Novel X – wt* | 0.25 |
| *Novel Y – wt* | $5.77e^{-03}$ |
| *Novel X – Novel Y* | 0.28 |
